# Reconstructing Ambient Temperature in Forensic Death Time Estimation

**DOI:** 10.1101/2025.11.14.688394

**Authors:** Jayant Shanmugam Subramaniam, Michael Hubig, Sebastian Schenkl, Holger Muggenthaler, Steffen Springer, Martin Weiser, Jakob Sudau, Faisal Shah, Gita Mall

## Abstract

Ambient temperature T_A_ has a strong impact on temperature based time since death estimation (TTDE). Frequently T_A_ is lowered instantaneously at a time t_0_ from a start value T_A0_ to T_A1_ < T_A0_ e.g. by window or door opening. We focus on reconstructing T_A0_ and t_0_.

TTDE literature suggests temperature measurements in closed compartments as e.g. cupboards or neighboring rooms, where T_A0_ could have been ‘preserved’ after t_0_. We aim to estimate t_0_ and T_A0_ from temperature measurements T_Z_(t) in closed compartments Z at times t > t_0_.

We obtained promising results assuming Newtonian cooling for boxes filled with air, with heaps of clothes or even with books in two different experimental scenarios. Two different parameter estimators, (t_0_^, T_A0_^) forsingle quadruple temperature measurement and (t_0_*, T_A0_*) for N quadruple measurements were tested.

Our results in a climate chamber were partially appropriate for TTDE input. A decline at time t_0_ from T_A0_ = 22.5°C ↓ T_A1_ = 14°C was reconstructed at t = t_0_ + 95min with relative deviations ρt_0_^ = 27% and ρT_A0_^ = 19% relative to t - t_0_ and T_A0_ – T_A1_ respectively, for N = 1 quadruple measurement with span Δt = 50min. For N = 200 quadruple measurements in [t = t_0_ + 95min, t = t_0_ + 295min] we found ρt_0_^ = 5% and ρT_A0_^ = 11% with the same Δt.

Further research is necessary to guarantee applicability in routine casework. We will investigate more elaborate cooling models, estimation algorithms, and evaluation localization.

## 1. Introduction

Though temperature based death time estimation (TTDE) is considered the most promising time of death estimation (TDE) method in forensic casework for short and medium times p.m., it is hampered by some serious and frequent drawbacks. Sudden unconsidered drastic changes of the ambient temperature T_A_ during body cooling (declines mostly) may lead to large errors in TTDE results (see [Weiser 2018], [Hubig 2010]). The changes in T_A_ may be caused by body transport as well as by opening doors or windows at the crime scene during the p.m. cooling phase before or after the body’s detection.

Since this problem is well known in forensic casework, there are heuristics in TTDE-literature (see [Henßge 1988], [Henßge 2002]) how to detect said T_A_-changes. As the problem is of some importance, the sources suggest T_A_-measurements in closed boxes or neighboring rooms of the crime scene to detect T_A_-changes during body cooling. It seems to be tempting to imagine the compartments could ‘preserve’ the original T_A_-value. This is merely an illusion as any compartment at the crime scene is externally exposed to a lowered ambient temperature T_A_ at time t > t_0_ and will begin to cool down from the original internal temperature T_A0_. This is supported by air exchange through small gaps in the compartments. In reality, one is confronted with the task of reconstructing T_A0_ from measurement results of T_Z_ ‘half way’ between T_A0_ and T_A1_.

Our approach for T_A0_-reconstruction is model based. Inspired by physics, we were looking for a thermodynamic model describing the cooling of the closed compartments contents, transforming the T_A0_-reconstruction task into a well-defined parameter estimation problem. Our cooling experiments with closed boxes and their contents in suddenly falling ambient temperature in a climate chamber suggest that small boxes with sufficient air content cool down approximately in a Newtonian process, described mathematically by a simple exponential curve. An additional complexity is the consequence of our need to not only estimate T_A0_ but the time t_0_ of the T_A0_-T_A1_-change as well. This induces a minimum number of two boxes with sufficiently different cooling curves and a minimum of one measurement quadruple with two measurements taken at each of the boxes.

## 2. Materials and Methods

### 2.1 Experiments

We performed two cooling experiments E1 and E2 in a climate chamber (3706/06 RMA 3313) of the company Feutron. The chamber guarantees a constant ambient temperature T_A_(t) as a function of time t in the limits +/-0.5°C. In every experiment two containers (a cupboard and a cardbox), called ‘boxes’ in the following, were placed in the climate chamber before the experiment was started. One box X contained air, while the other box Y contained books in experiment E1 and a heap of clothes in the other experiment E2. Both of the boxes X and Y were closed but not airtight, simulating closed compartments like cupboards at a crime scene. All of our measurement equipment was from the series ALMEMO of the company AHLBORN. We placed a temperature probe (FVAD 05-TOK300) in a box X to measure the air temperature T_X_. Another sensor (FN0001K NTC-Sensor) was placed below the material in box Y to measure the temperature T_Y_ on the bottom of the box. Moreover, another temperature probe (FVAD 05-TOK300) was placed in the chamber next to the boxes X and Y to register the chambers ambient temperature T_A_. The temperatures T_A_, T_X_, T_Y_ were registered during the whole experiment every minute using datalog devices (MA7120 and MA809).

Each experiment comprised two successive time intervals P0 and P1 and started in interval P0. At a certain point in time t_0_ the interval P0 ended and interval P1 began and lasted until the end of the experiment. During interval P0 the chamber was kept at a constant ambient temperature T_A0_. At time t_0_ the temperature was instantaneously downregulated from T_A0_ to a temperature T_A1_ and kept constant during interval P1.

Fig. 1 shows the experiment E1’s setup where Y was a cardbox filled with books.

**Fig. 1:**
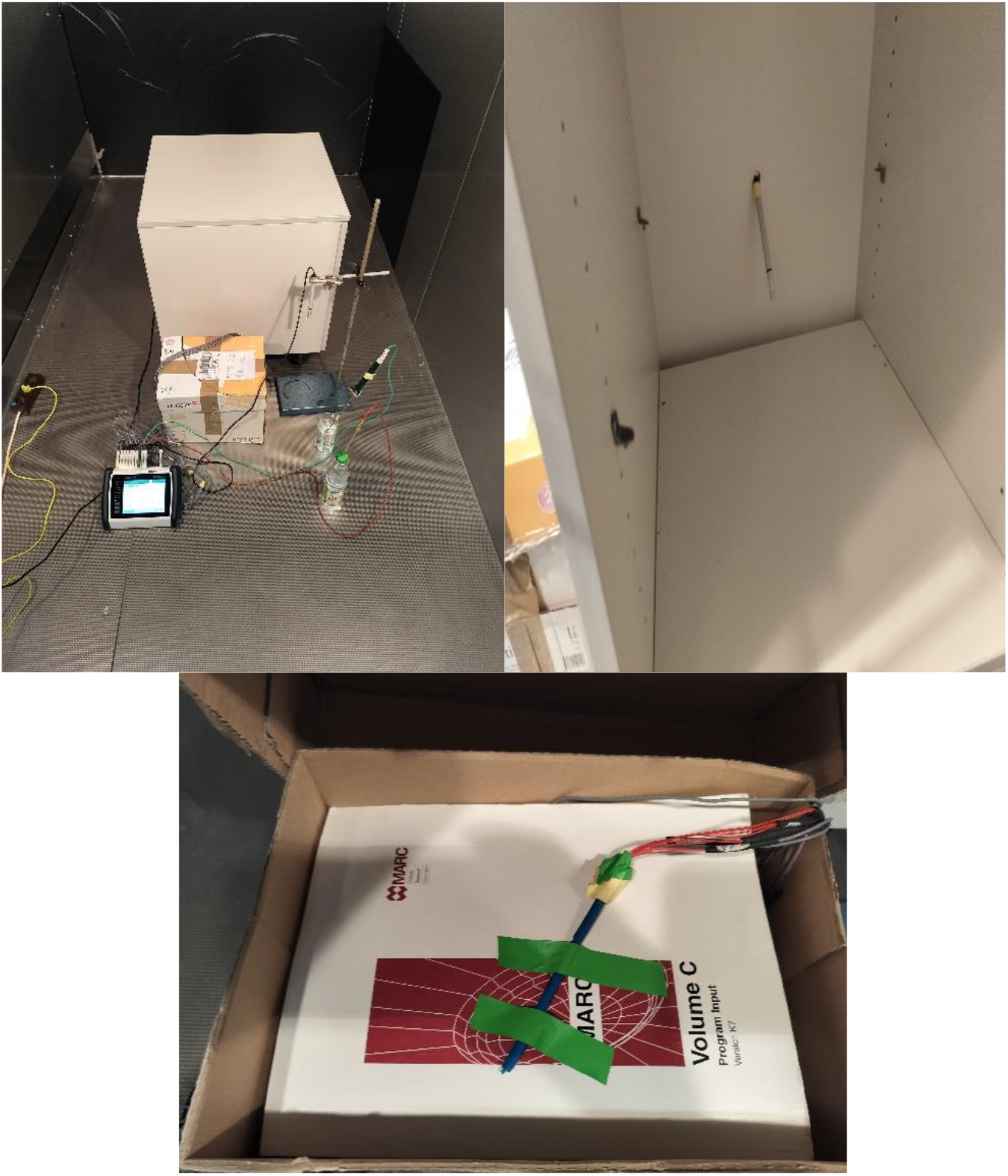
Experiment E1: *Upper left:* Scenario in climate chamber (**1a:** upper left) with X: cupboard (background), Y: cardbox with books (middle, left), ambient probe for T_A_ (middle right), datalogger (foreground left) temperature probes in water bottles (not used for evaluation) (foreground right). **1b:** *upper right:* Interior of cupboard X with probe for T_X_. **1c:** *lower:* Interior of cardboard Y with books and probe for T_Y_.

### 2.2 Estimate of t_0_ and T_A0_ from single quadruple measurement of temperatures in boxes X and Y via Newtonian cooling

The following abbreviations and symbols are used:

T_A0_ = Initial ambient temperature (unknown)

T_A1_ = Constant ambient temperature during cooling (measured)

T_X_(t) = Temperature of content at time t in a closed box X (measured)

T_Y_(t) = Temperature of content at time t in a closed box Y (measured)

T_Z,1_ = Temperature of content at time t_1_ in a closed box Z = X, Y (measured)

T_Z,2_ = Temperature of content at time t_2_ in a closed box Z = X, Y (measured)

t_0_ = Time of initial instantaneous temperature decline:

T_A0_ (unknown) ==> T_A1_ (measured)

t_1_ = Time of first two temperature measurements T_X,1_, T_Y_,_1_ during cooling

t_2_ = Time of second two temperature measurements T_X,2_, T_Y,2_ during cooling Δt = Time difference between t_1_ and t_2_ called *quadruple span*

The measurements T_X_,_1_, T_X_,_2_, T_Y_,_1_, T_Y_,_2_ constitute one *quadruple* of temperature measurements. It is convenient though not necessary to choose two measurement times t_1_ and t_2_ only and to perform synchronous measurements T_X_,_1_, T_Y_,_1_ and T_X_,_2_, T_Y_,_2_ respectively at each one of the points t_1_ and t_2_ in time.

Newtonian cooling for T_Z_ = T_X_, T_Y_ at time t is given by

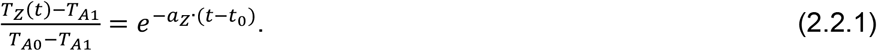

Taking the logarithm of (2.2.1) yields

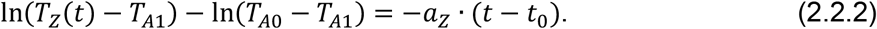

Inserting T_Z_ = T_X,1_, T_X,2_ and t = t_1_, t_2_ into (2.2.2) and subtracting the two resulting equations

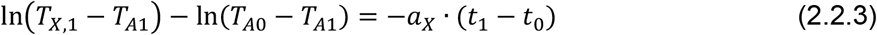

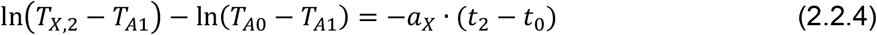

yields

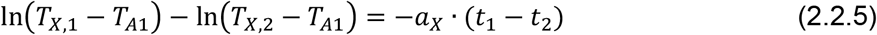

and thus the factor a_X_:

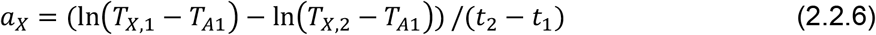

Analogously, *a*_*Y*_ is obtained from T_Z_ = T_Y,1_, T_Y,2_.

With known *a*_*X*_, *a*_*Y*_, inserting *T*_*Z*_ = *T*_*X*,1_, *T*_*Y*,1_ into (2.2.2) results in

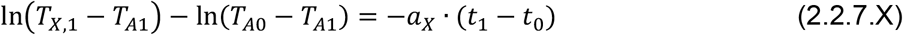

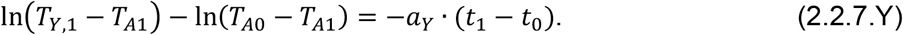

Subtracting (2.2.7.Y) from (2.2.7.X) yields

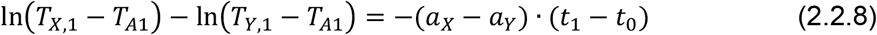

and thus an estimate for t_0_:

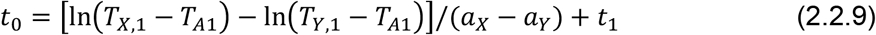

Finally, solving (2.2.1) for T_A0_ with *T*_*Z*_ = *T*_*X*,1_ yields

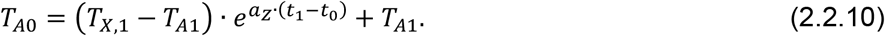

Now, (2.2.6) can be used to estimate a_Z_, while (2.2.9) and (2.2.10) give single quadruple estimators 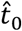 and 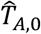 for the unknown parameters t_0_ and T_A0_.

### 2.3 Estimate from n quadruple measurements of temperature T in boxes X and Y via Newtonian cooling

Let T_A0_,T_A1_, t_0_ be as in 2.2, let further for all n = 1,…, N be t_1,n_ and t_2,n_ be the time points of temperature measurements T_X,1,n_, T_Y,1,n_ and T_X,2,n_, T_Y,2,n_ in the boxes X and Y respectively with a time distance Δt = t_2,n_ - t_1,n_ independent of n. The difference between the first measurement times t_1,n_ and t_1,(n+1)_ of two consecutive quadruples is dt = t_1,(n+1)_ – t_1,n_, which is chosen independently of n as well. Thus we have N = (t_1,N_ – t_1,1_) / dt +1.

The single estimators t_0,n_^ and T_A0,n_^ of the time t_0_ since temperature drop and the ambient temperature T_A0_ before the drop at t_0_ are then computed as in 2.2:

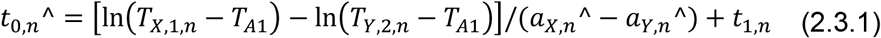

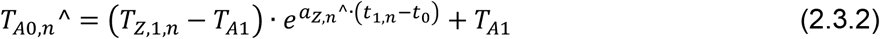

The Weighted Mean Estimators (WME) t_0_* and T_A0_* are yielded by computing the weighted mean of all of the respective single estimators t_0,n_^ and T_A0,n_^ from the N equations in (2.3.1) and in (2.3.2) respectively with the weight factors W_1_, …, W_N_ for the N equations in (2.3.1) and for (2.3.2). Weighting of the mean computation with the factors W_n_ gives the WMEs t_0,n_* and T_A0,n_*:

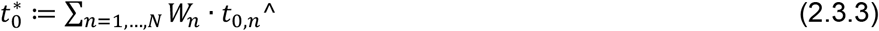

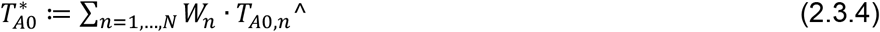

Choosing the weights W_n_ inversely proportional to the standard deviations S(t_0,n_^) and S(T_A0,n_^) of the single estimators t_0,n_^ and T_A0,n_^ in the equations (2.3.1) and (2.3.2) respectively is the canonical approach:

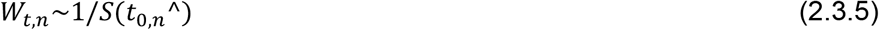

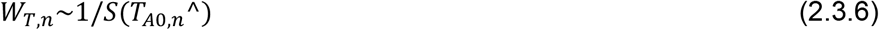

with:

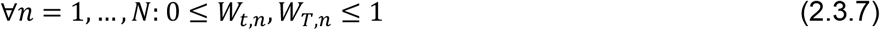

and:

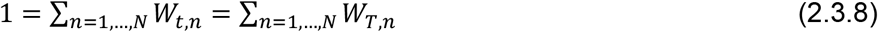

The time difference between the measurement times t_1,n_ and t_1,(n+1)_ of two consecutive quadruples with indices n and n+1 is named dt and *will be kept constant in all of our experiments and computations*:

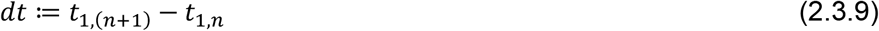

### 2.4 Estimate weights and errors of the method

The usual Gaussian error propagation approach – estimating standard deviations via linear approximation of random variables - yielded the standard deviation estimators St_0_^ and ST_A0_^ for the single quadruple approach as well as St_0_* and ST_A0_* for the WMEs with multiple quadruple measurements. The standard deviation estimations were used for error quantification as well as for weight computation in the WME. We assumed the input random variables T_A1_, T_X,1_, T_X,2_, T_Y,1_, T_Y,2_, t_1_, t_2_ to be independently and identically normally distributed with standard deviation ST = 0.1°C for temperature variables T and St = 1min for time variables t. Denoting the true values by t_0_ ^+^ and T_A0_^+^, and the estimated values by t_0_ ^#^ ( = t ^ or t *) and T ^#^ ( = T ^ or T *), we can write the errors *δ*t and *δ*T as

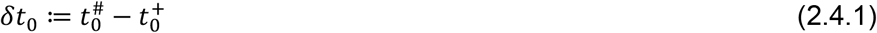

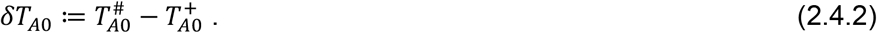

With fixed values for time difference Δt = t_2_ – t_1_ the relative errors ρt_0_(t_1_), ρT_A0_(t_1_) were computed as

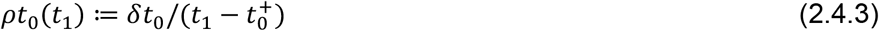

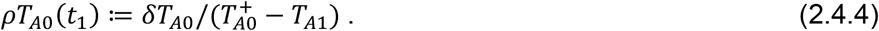

The relative dispersions Rt_0_(t_1_), RT_A0_(t_1_) or relative standard deviations were computed from the standard deviations St_0_ ^#^(t), ST ^#^(t) as

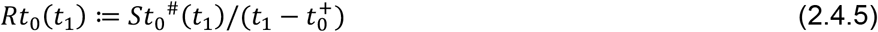

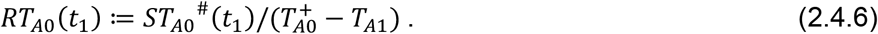

## 3. Results

The parameters for the experiments E1 and E2 were chosen as follows:

- E1: t_0_ = 305.00min; T_A0_ = 22.50°C; T_A1_ = 14.00°C
- E2: t_0_ = 158.13min; T_A0_ = 22.16°C; T_A1_ = 8.71°C

For the tests on experiment E2 data, the temperature T_A1_ of the lower plateau was estimated using the mean of the measured values T_A_(t) for all t ≥ t_01_ for a fixed threshold t_01_ := 180min, which was the earliest value t_1_ we considered, whereas the temperature plateau of time interval P0 ended at t = 150min. The measured ambient temperature T_A_(t) did not fall instantaneously from T_A0_ to a constant level T_A1_ < T_A0_, but needed about 20 min to reach its overshot-minimum 7°C and subsequently climbed asymptotically towards a constant level which, however, was not reached during measurement time (see Fig.2 lower part). The mean estimated temperature in P0 computed from T_A_(t) values with t ≤ 150min was T_A0_ = 22.16°C, while the mean estimated temperature in P1 computed from T_A_(t) values with t ≥ 180min was T_A1_ = 8.71°C.

**Fig. 2:**
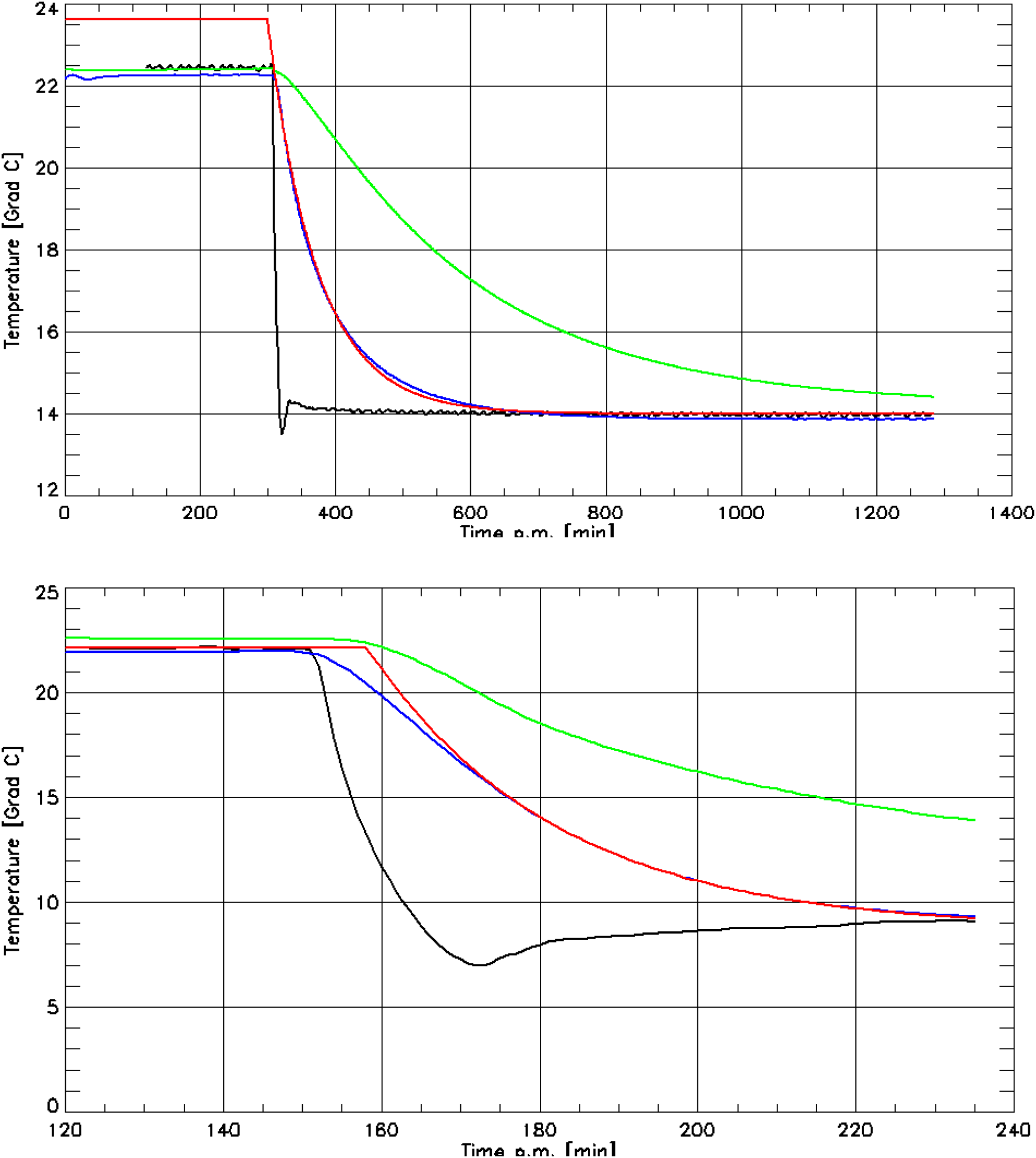
(**2a:** *upper part:)* Experiment E1, (**2b:** *lower part:)* Experiment E2. Measured curves: ambient temperature T_A_(t) (black), air temperature T_X_(t) (blue) in box X, temperature T_Y_(t) (green) on the bottom of box Y below materials (E1: books, E2: pile of clothes). The reconstructed cooling curve of T_X_^ (red) demonstrates the estimated values T_A0_* and t_0_* for single measurement quadruple estimation. The measurement times were t_1_ = 400min, t_2_ = 650min (Δt = 250min) for E1 and t1 = 180min and t_2_ = 190min (Δt = 10min) for E2 respectively.

Though the E2 data did not exactly match our ideal scenario of two ambient temperature plateaus at time-constant temperatures T_A0_ and T_A1_ respectively, we kept the E2 data to test the robustness of our approach against deviations from this scenario, frequently met in application cases.

### 3.1 Evaluation with one measurement quadruple

Fig.2 consists of two example diagrams showing the measured temperature curves T_A_(t) (black), T_X_(t) (blue), T_Y_(t) (green) for the experiments E1 and E2 respectively, as well as the curve T_X_^(t) (red), computed via (2.2.10) using the parameters estimation results t_0_^ and T_A0_^ respectively. For E1, the times t_1_, t_2_ for the respective two measurements were t_1_ = 400min, t_2_ = 650min, whereas for E2 we chose t_1_ = 180min and t_2_ = 190min. Note that the black, blue and green curves were identical for all measurement times t_1_, t_2_ in each of the experiments E1, E2.

The results of all one-quadruple estimations are represented in Tab.1 for E1 and in Tab.2 for E2.

**Tab 1:**
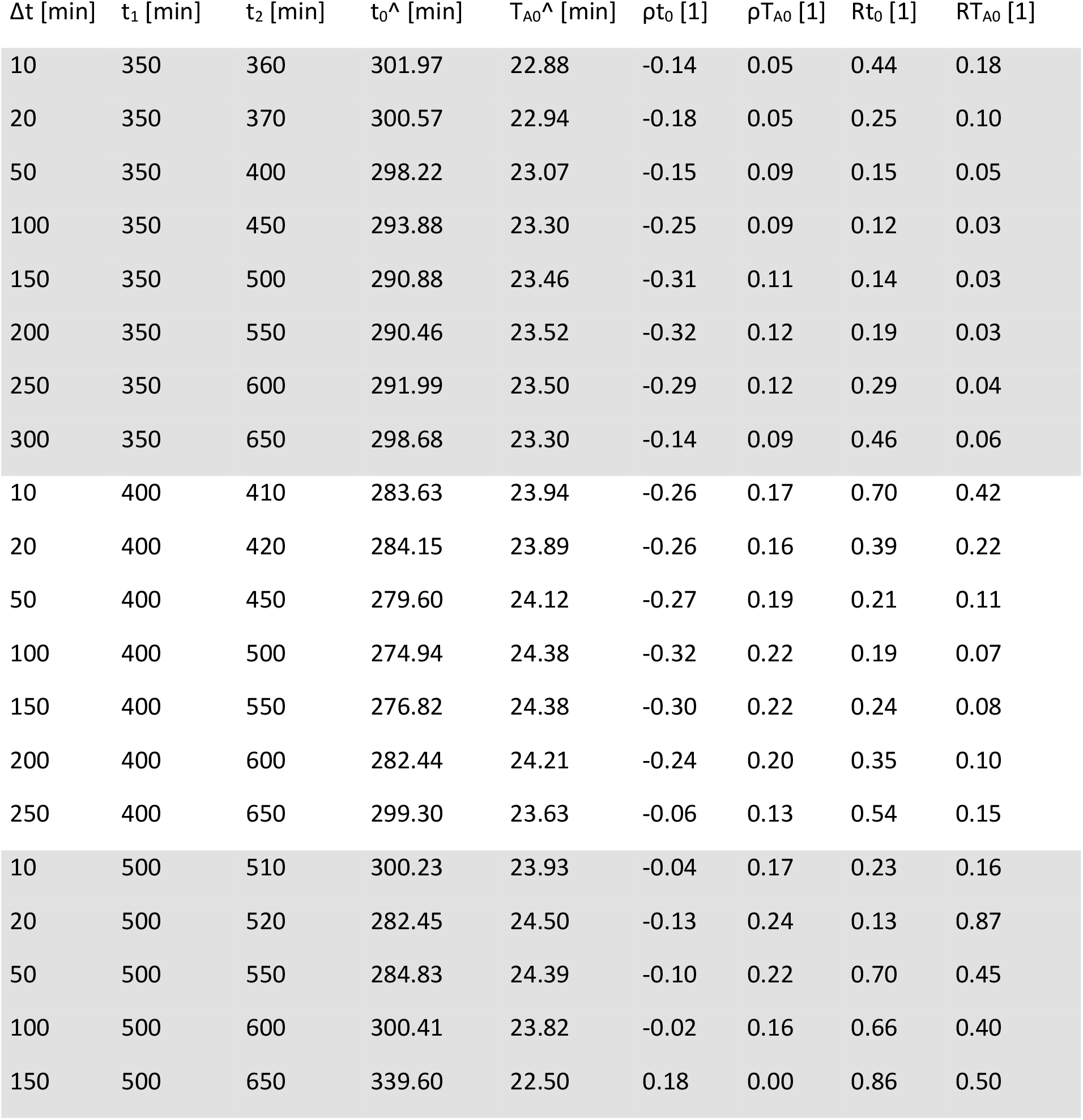
Results of estimations in E1 for only one measurement quadruple. Quadruple span Δt, first measurement time t_1_ and last measurement time of the first quadruple; estimation values t_0_^ for time limit t_0_ and T_A0_^ for temperature T_A0_ of first interval P0; errors *δ*t_0_ and *δ*T_A0_ as well as relative errors ρt_0_ and ρT_A0_ and relative standard deviations Rt_0_ and RT_A0_ of the estimators.

**Tab 2:**
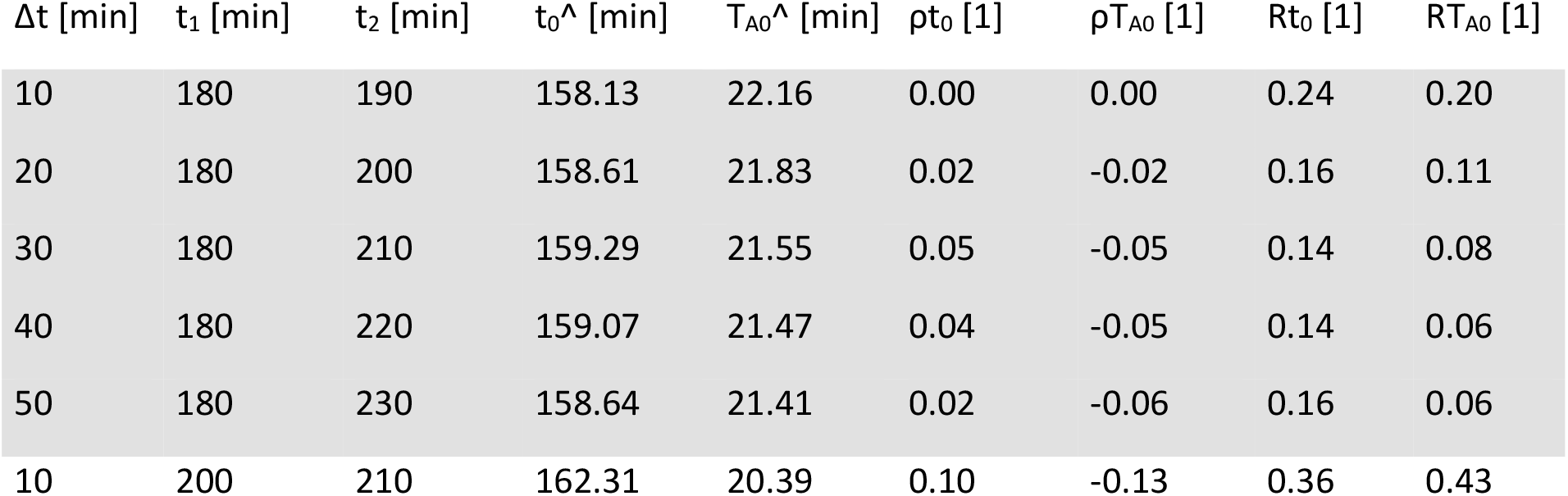
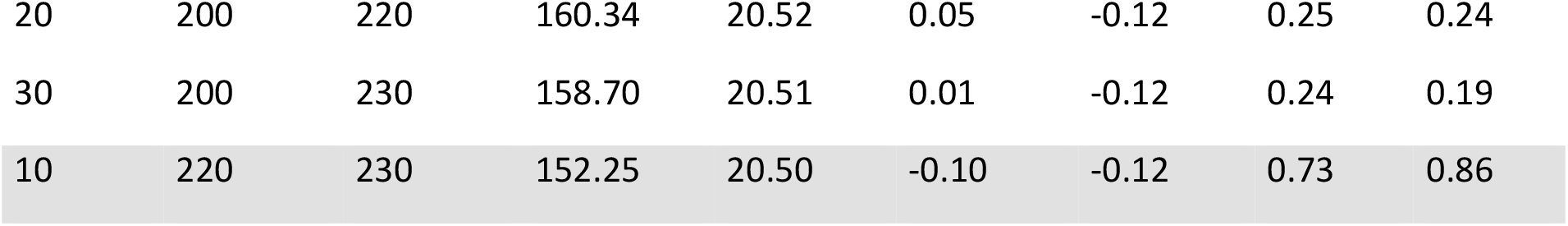
Results of estimations in E2 for only one measurement quadruple: Quadruple span Δt, first measurement time t_1_ and last measurement time of the first quadruple; estimation values t_0_^ for time limit t_0_ and T_A0_^ for temperature T_A0_ of first interval P0; errors *δ*t_0_ and *δ*T_A0_ as well as relative errors ρt_0_ and ρT_A0_ and relative standard deviations Rt_0_ and RT_A0_ of the estimators.

### 3.3 Error estimation for one quadruple estimation

In Fig.3 the relative standard deviations Rt_0_(t) and RT_A0_(t_1_) are shown as functions of the first measurement time t_1_. The diagrams present the curves for fixed time differences Δt = 10min, 20min, 30min, 40min, 50min in experiment E1 and for Δt = 10min, 20min, 30min, 40min, 100min in experiment E2. The coefficients a_X_ and a_Y_ were set to:

- a_X_ = 0.01288min^-1^, a_Y_ = 0.00298min^-1^ for E1 yielded in paragraph 3.1 with t_1_ = 350min, t_2_ = 400min.
- a_X_ = 0.04204min^-1^, a_Y_ = 0.01428min^-1^ for E2 yielded in paragraph 3.1 with t_1_ = 180min, t_2_ = 190min.

### 3.3 Evaluation with several measurement quadruples

The evaluation with several measurement quadruples were carried out with varying times of first quadruple measurement t_1,1_ and varying last quadruple measurement time t_1,N_. As the time interval between quadruples, dt = 1min, remained constant, the resulting number N of quadruples per evaluation varied accordingly.

Fig.5 presents an example of estimation results for approaches with several measurement quadruples: The two temperature-time diagrams present the measured curves T_A_(t) (black), T_X_(t) (blue), T_Y_(t) (green) for the experiments E1 and E2 respectively. The curves T_X_*(t) (red) and T_Y_*(t) (red) were reconstructed using the estimator values a_X_* and a_Y_*, which are interim results of computing the final estimation results t_0_* and T_A0_*. For E1 the first quadruple’s first measurement time was t_1,1_ = 500min (pink mark left) while the last quadruple’s first measurement time was t_1,N_ = 599min (pink mark right) which adds up to N = 100 quadruples. For E2 the number of measurement quadruples was N = 20 with the first measurement time t_1,1_ = 180min (pink mark left) of the first quadruple and the last quadruples first measurement at t_1,N_ = 199min (pink mark right). The time difference Δt = t_2,n_ – t_1,n_ was chosen Δt = 20min for all quadruples, while the time distance dt = t_1,(n+1)_ – t_1,n_ between the first measurements of two successive quadruples was for all evaluations dt = 1min.

The results of all N > 1 quadruple-estimations are represented in Tab.3 for E1 and in Tab.4 for E2.

**Tab 3:**
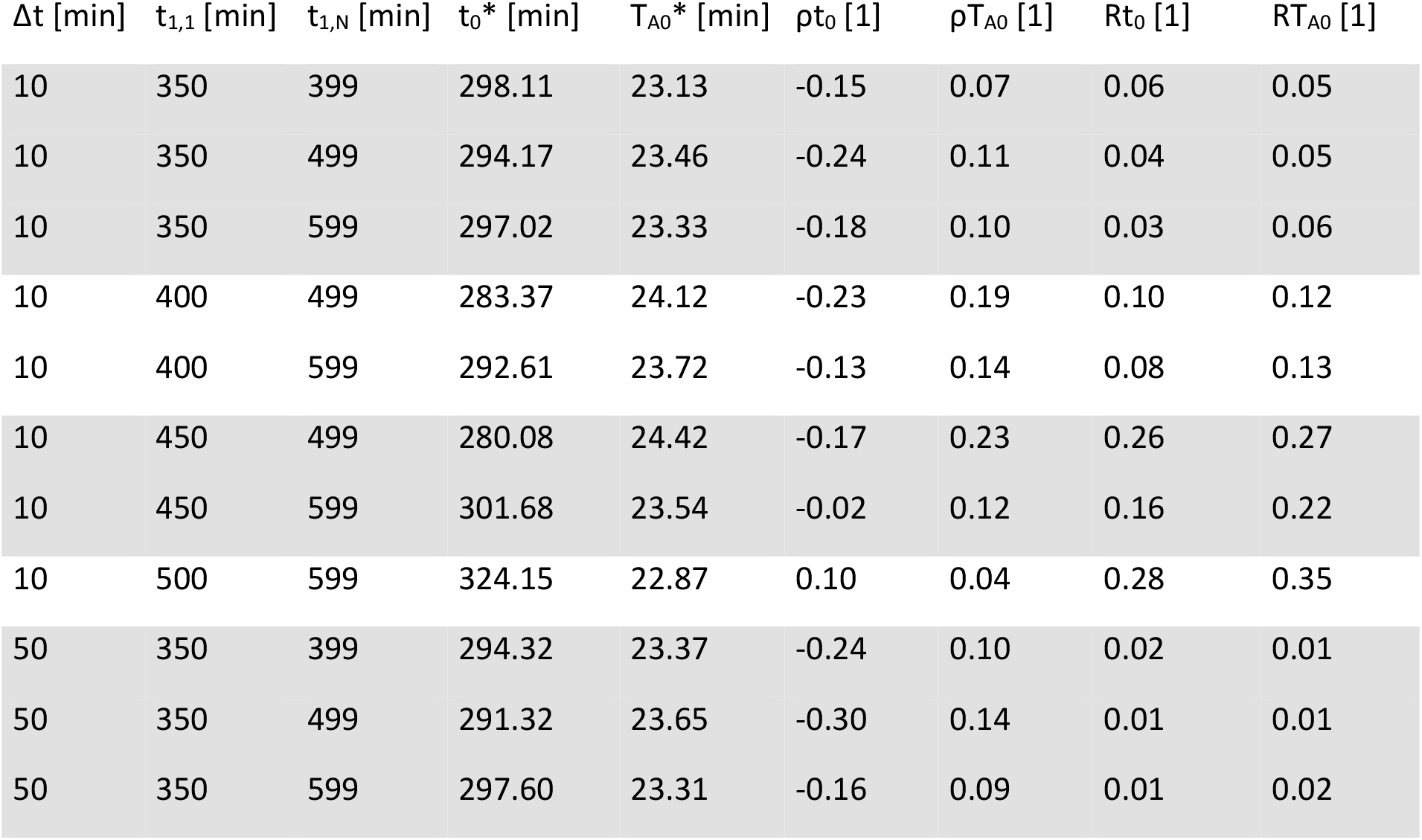
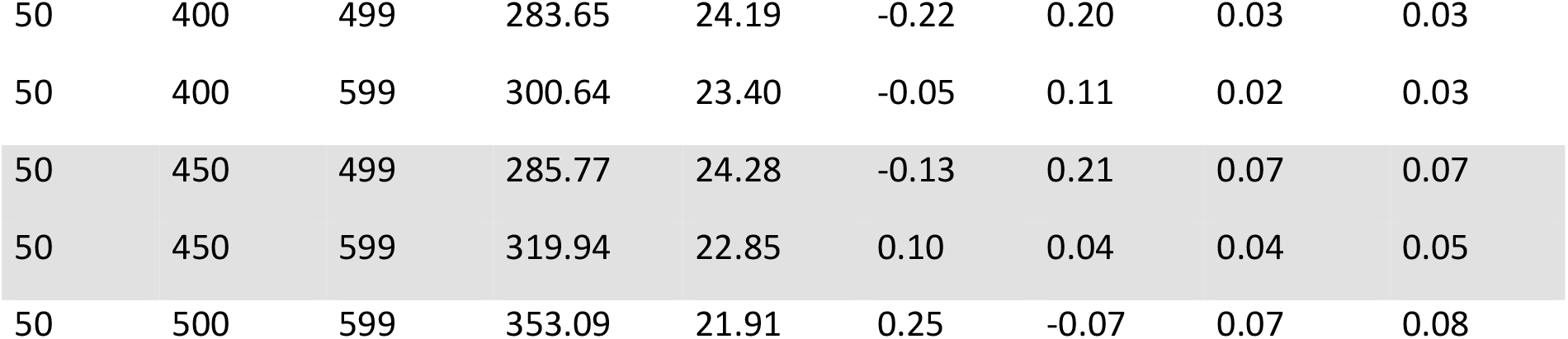
Results of estimations in E1 for N > 1 measurement quadruples. Quadruple span Δt, first measurement time t_1,1_ of first quadruple and first measurement time t_1,N_ of the last quadruple; estimation values t_0_* for time limit t_0_ and T_A0_* for temperature T_A0_. Relative errors ρt_0_ and ρT_A0_ and relative standard deviations Rt_0_ and RT_A0_ of the estimators. The distance between neighbor-quadrupels is dt = 1min and the number N = (t_1,N_ – t_1,1_) / dt + 1 of quadruples varies accordingly.

**Tab 4:**
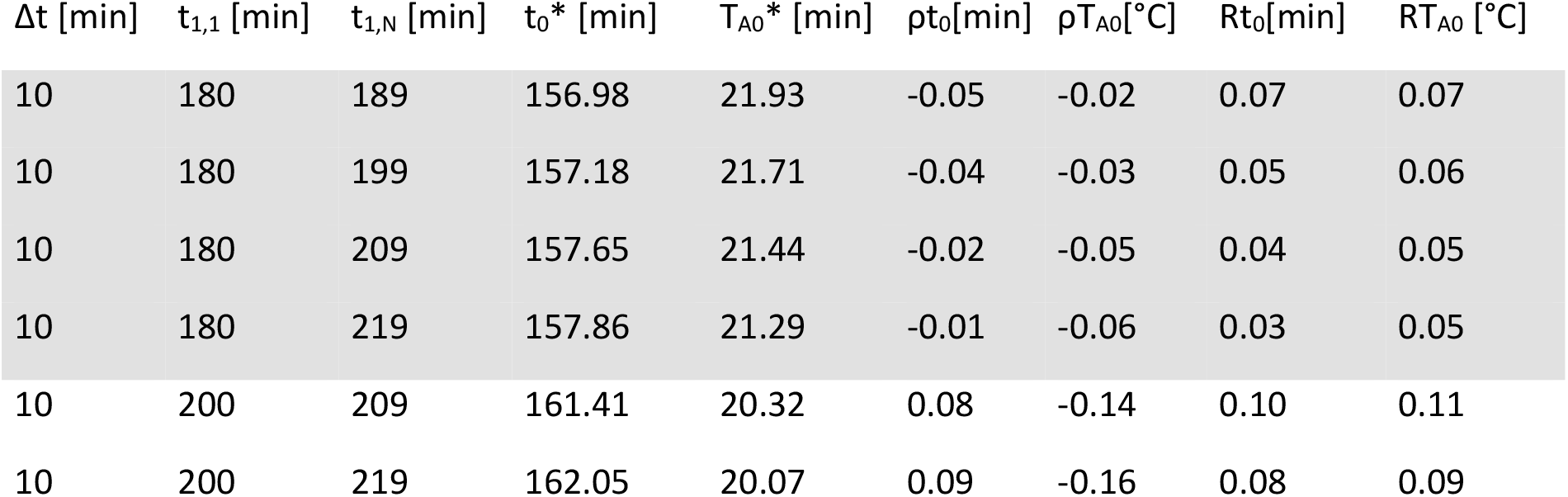
Results of estimations in E2 for N > 1 measurement quadruples. Quadruple span Δt, first measurement time t_1,1_ of quadruple 1 and first measurement time t_1,N_ of the last quadruple; estimation values t_0_* for time limit t_0_ and T_A0_* for temperature T_A0_. Relative errors ρt_0_ and ρT_A0_ and relative standard deviations Rt_0_ and RT_A0_ of the estimators. The distance between neighbor-quadrupels is dt = 1min and the number N = (t_1,N_ – t_1,1_) / dt + 1 of quadruples varies accordingly.

## 4. Discussion

Errors in ambient temperature T_A_ are of extreme importance in TTDE (see e.g. [Weiser 2018], [Hubig 2010]). There have been several attempts to deal with – or even to overcome this problem. In practical TTDE casework, there are frequent scenarios, where a constant indoor ambient temperature is significantly reduced by scene manipulations (see [Sauer 2024]). The latter sudden temperature drops typically occur during indoor homicide investigations when crime scene investigators, unaware of the thermodynamical consequences, open windows for ventilation before ambient temperature measuring takes place.

One approach [Mall 2005] to deal with said situations showed under the assumption of the Marshall & Hoare model with parameter determination of Henßge (MHH) that the information about a sudden T_A_-decline during body cooling is contained in the cooling curve itself and can in principle be extracted numerically. The Prism method [Potente 2021] aims for the same goal using pairs of temperature-time measurements to estimate MHH parameters. Estimating the temperature drop from body cooling alone is, however, a severely ill-conditioned problem and thus subject to large errors introduced by unavoidable measurement noise. Including additional external information is the best way to improve the reconstruction accuracy. As mentioned in the introduction we use temperature measurements in closed compartments at the crime scene. We reconstruct the initial ambient temperature T_A0_ after one sudden temperature decline down to a temperature plateau at T_A1_ at the time t_0_ after the beginning of body cooling.

The estimated values of t_0_ and T_A0_ can serve as input for TTDE using a Finite Element Model (FEM) as in [Shanmugam 2023], [Ullrich 2023].

At first glance, our experimental approach may appear overly complex: Would it not be simpler to use just one box X? Given that three parameters – a_X_, t_0_, T_A0_ – are to be estimated, would it not suffice to perform three temperature measurements T_1_, T_2_, T_3_ at three different times t_1_, t_2_, t_3_ in this box? By substituting our measurement results into the model formula (2.2.2), we would obtain three equations and attempt to solve for the unknown parameters. However, this approach is hindered by the fact that a single box with a single cooling rate a_X_ yields only two independent equations – one equation short of what is required to determine three parameters. Hence, it is necessary to use a second box Y with a different cooling rate a_Y_ ≠ a_X_.

There were conditions where a sudden decline at time t_0_ from T_A0_ = 23°C to T_A1_ = 10°C of the ambient temperature in a climate chamber could be reconstructed at t = t_0_ + 95min with relative deviations ρt_0_^ = 27% and ρT_A0_^ = 19% of the estimators based on N = 1 quadruple measurement with a span of Δt = 50min. In case of N = 200 quadruple measurements starting at t = t_0_ + 95min and ending at t = t_0_ + 295min we found for weighted mean estimators distinctively reduced relative deviations ρt_0_^ = 5% and ρT_A0_^ = 11% with the same quadruple span Δt = 50min. Relative errors ρt_0_ = 10% and ρT_A0_ = 4% were obtained from a quadruple span Δt = 50min with t_1,1_ = 450min and t_1,N_ = 599min (N = 150).

Clearly the usage of N > 1 quadruple measurements seems to improve reconstruction quality in comparison to the single quadruple approach N = 1: Comparison of Fig.3 and Tab.3 show, e.g., for N = 100, t_1,1_ = 400min, t_1,1_ = 499min, Δt = 10min the relative standard deviations Rt_0_(t_1,1_) = 10%, RT_A0_(t_1,1_) = 12%, whereas in case N = 1 we have Rt_0_(t_1_) = 70%, RT_A0_(t_1_) = 40% for t_1_ = 400min. This trend can be detected in Tab.3 and Tab.4 as well. We observe generally decreasing values of Rt_0_(t_1,1_) and RT_A0_(t_1,1_) with constant t_1,1_ and rising t_2,1_ (which means rising N due to constant dt = 1min) at constant levels of the quadruple span Δt. In Tab.4 we have e.g. for E2 with Δt = 10min and t_1,1_ = 180min for t_1,N_ rising from t_1,N_ = 189min up to t_1,N_ = 219min (which means N rising from N = 10 to N = 30) decreasing relative standard deviations from Rt_0_(t_1,1_) = 7% to Rt_0_(t_1,1_) = 3% and from RT_A0_(t_1,1_) = 7% to RT_A0_(t_1,1_) = 5% respectively.

Increasing values of t_1_ or t_1,1_, which means later measurement start times, generally lead to larger values of Rt_0_ and RT_A0_: For N = 1 this can be seen in the diagrams of Fig. 3 for E1: All RT_A0_(t_1_) curves and for t_1_ > 340min (for E1) and t_1_ > 170min (for E2) all Rt_0_(t_1_) curves are monotonically increasing with measurement begin t_1_. For N > 1 we see in Tab.3 for E1 monotonically increasing values of Rt_0_(t_1,1_) and RT_A0_(t_1,1_) for increasing t_1,1_ (e.g. for Δt = 10min, t_1,1_ = 400min ↑ 500min and t_1,N_ = 499min ↑ 599min (N = 100): Rt_0_(t_1,1_) = 10% ↑ 28% and RT_A0_(t_1,1_) = 12% ↑ 35%), as well as in Tab.4 for E2 (e.g. for Δt = 10min, t_1,1_ = 180min ↑ 200min, t_1,N_ = 189min ↑ 209min (N = 10): Rt_0_(t_1,1_) = 7% ↑ 10% and RT_A0_(t_1,1_) = 7% ↑ 11%). Thus we notice that larger time intervals between the first and the last quadruple measurement are generally preferable.

**Fig. 3:**
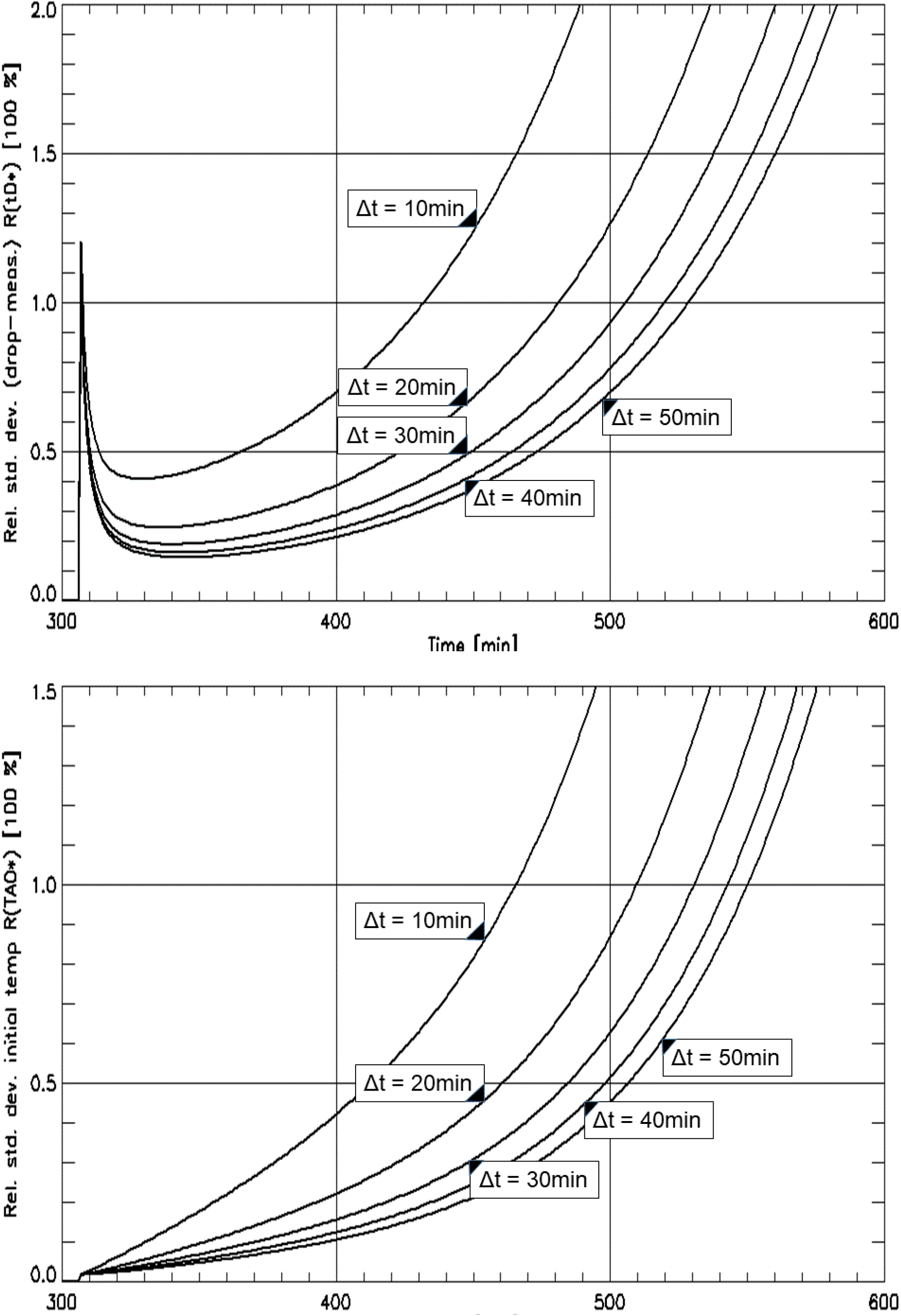
For experiment E1: Relative standard deviations Rt_0_(t_1_) (in **3a:** *upper part*) and RT_A0_(t_1_) (in **3b:** *lower part*) as a function of the first measurement position t_1_ for different distances Δt = t_2_ – t_1_ as given in the labels.

The influence of the quadruple span Δt = t_2_ – t_1_, or Δt = t_2,n_ – t_1,n_ respectively, in experiment E1 (see Fig.3) is clear as well: For single quadruples (N = 1) the relative standard deviations Rt_0_(t_1_) and RT_A0_(t_1_) are consistently decreasing with increasing span Δt. We assume that this dependency is disturbed in experiment E2 (see Fig.4) due to the experimental deviation of T_A_(t) from our model assumption of two adjacent plateaus with constant temperatures T_A0_ and T_A1_ respectively and an instantaneous descent at t_0_ between them. For multiple quadruple measurements (N > 1) Tab.3 shows the same trend: E.g. for Δt = 10min and t_1,1_ = 350min and t_1,N_ = 399min (which means N = 50) we have Rt_0_(t_1,1_) = 6% and RT_A0_(t_1,1_) = 5%, whereas Δt = 50min and t_1,1_ = 350min and t_1,N_ = 399min gives lower values Rt_0_(t_1,1_) = 2% and RT_A0_(t_1,1_) = 1%. It is important to note that the standard deviations S(t_0_) and S(T_A0_) required for Rt_0_ and RT_A0_ were computed using Gaussian error propagation, assuming compatibility of the Newtonian cooling model. Therefore the relative errors ρt_0_ and ρT_A0_ are more suitable indicators of model quality than the relative standard deviations Rt_0_(t_1,1_) and RT_A0_.

**Fig. 4:**
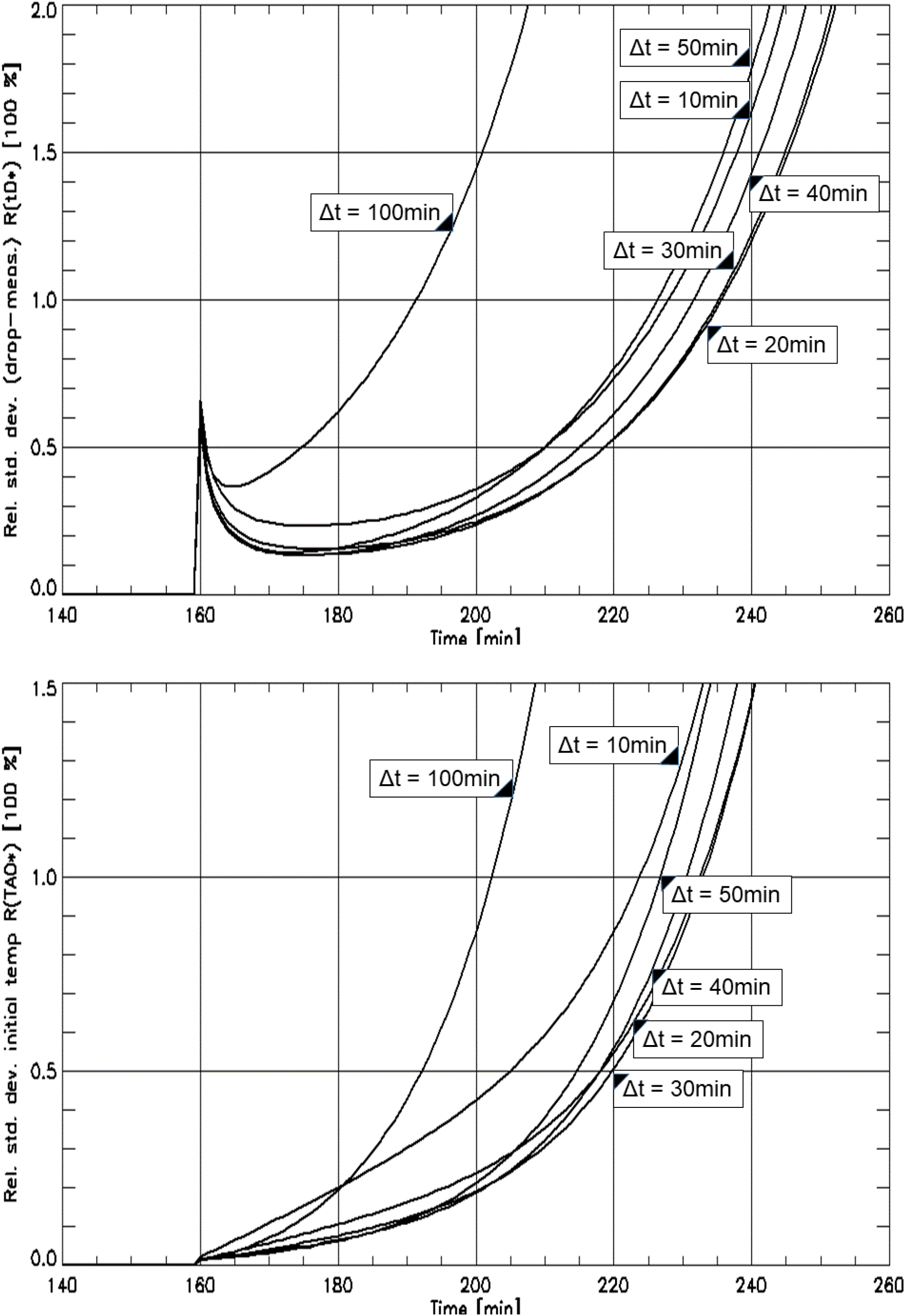
For experiment E2: Relative standard deviations Rt_0_(t_1_) (in **4a:** *upper part*) and RT_A0_(t_1_) (in **4b:** *lower part*) as a function of the first measurement position t_1_ for different distances Δt = t_2_ – t_1_ as given in the labels.

**Fig. 5:**
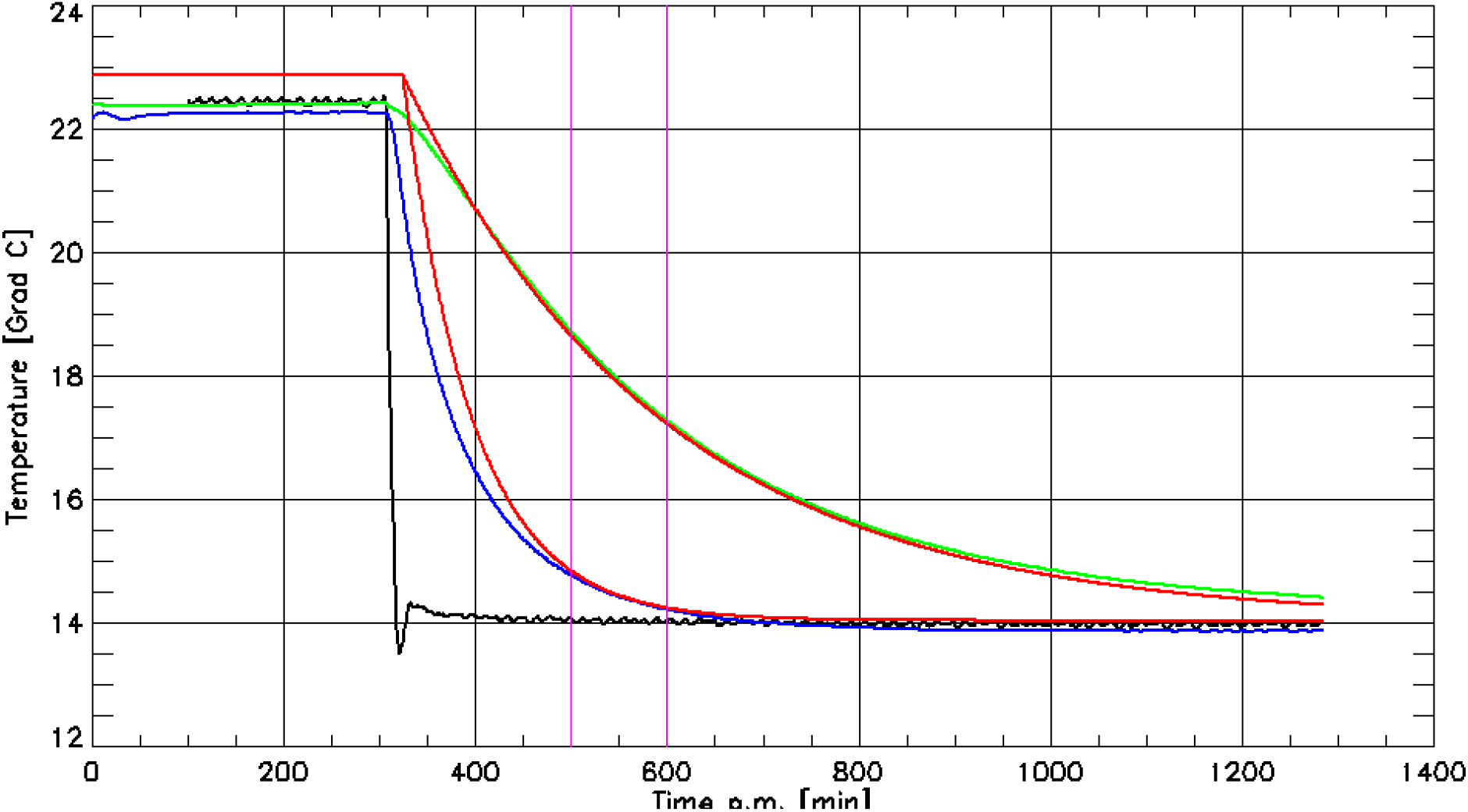
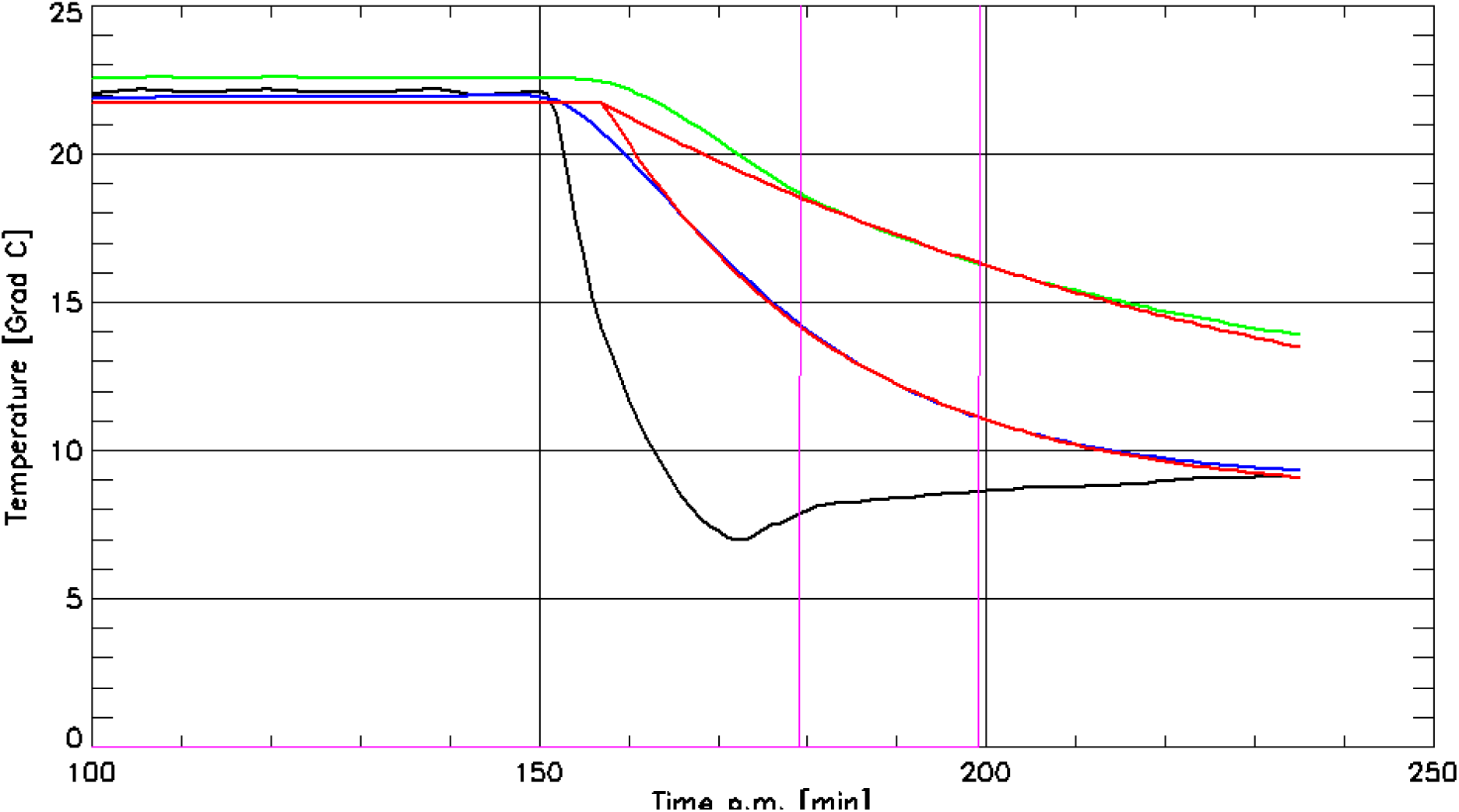
In **5a:** *upper Part:* Experiment E1, in **5b**: *lower part:* Experiment E2. Measured curves: ambient temperature T_A_(t) (black), air temperature T_X_(t) (blue) in box X, temperature T_Y_(t) (green) on the bottom of box Y below materials (E1: books, E2: pile of clothes). The reconstructed cooling curves of T_X_* and T_Y_* (red) demonstrate the estimated values T_A0_* and t_0_* for multiple measurement quadruple estimation. The points in time t_1,1_ and t_1,N_ of measurement begin for the first and for the last quadruple are marked by thin rectangular lines (pink). The distance between neighbor-quadrupels is dt = 1min and the number N = (t_1,N_ – t_1,1_) / dt + 1 of quadruples varies accordingly.

Looking at the signs of the N = 1 quadruple estimator’s relative errors ρt_0_ and ρT_A0_ we see nearly constant negatives in case of t_0_ and a nearly constant positive relative error for T_A0_ in Tab.1 for E1. This is reversed in Tab.2 for E2. Both biases are most probably the result of our (strongly simplifying) Newtonian cooling model assumption. The discrepancy between the relative errors ρt_0_ and ρT_A0_ compared to the relative standard deviations Rt_0_ and RT_A0_ in Tab.3 and Tab.4 appears to stem from our simplified model assumption as well. Assuming the error distributions to be Gaussians, we would expect ca 68% of the relative errors ρt_0_(t_1_) and ρT_A0_(t_1_) to fall into the intervals It_0_ := [-Rt_0_(t_1_), Rt_0_(t_1_)] and IT_A0_ := [-RT_A0_(t_1_), RT_A0_(t_1_)] respectively. However, this expectation is not met in all cases. In Tab.3 for Δt = 10 and t_1,1_ = 450 and t_1,1_ = 500 we have ρt_0_ in It_0_ and ρT_A0_ in IT_A0_ and in Tab.4 only three out of six cases satisfy the condition.

Comparing relative errors ρt_0_ and ρT_A0_ in Tab.1 and Tab.3 again suggests that Newtonian cooling might be an overly simplistic model. For instance, the single quadruple N = 1 measurement determination for Δt = 10min and t_1_ = 350min shows relative errors ρt_0_ = −14% and ρT_A0_ = 5% whereas the determination with N = 100 quadruples for Δt = 10min, t_1,1_ = 350min and t_1,N_ = 499min leads to even slightly higher relative errors ρt_0_ = −15% and ρT_A0_ = 7%. Examining the measured - and the fitted curve with N = 100 shows a very good fit in the interval [t_1,1_, t_1,N_ + Δt] of the quadruple measurements, however notable errors in t_0_ and T_A0_ are still present. This seems to be a perfect case of ‘underfitting’: Despite optimum matching in the area used, the model is not able to correctly reconstruct the curve shape outside the input measurement interval. This flaw is not resolved by enlarging the fitting interval [t_1,1_, t_1,N_ + Δt] which means rising N equivalently: For Δt = 10min, t_1_ = 350min and t_1,N_ = 599min (N = 250) the relative errors increased slightly to ρt_0_ = −18% and ρT_A0_ = 10%. Again the graphs of the measured and reconstructed curves T_X_(t), T_Y_(t), T_X_*(t), T_Y_*(t) presented correct matching in [t_1,1_, t_1,N_ + Δt] but nevertheless the errors ρt_0_ and ρT_A0_ persisted.

The measurements and computations presented were performed in a laboratory setting under maximum control of the relevant variables. In practical casework additional challenges will arise such as deviations of the ambient temperature curve T_A_(t) from our instantaneous drop from a constant plateau at T_A0_ down to another constant plateau at T_A1_ << T_A0_. In our experiment E2 significant deviations of the measured curve T_A_(t) from the two-plateau-scenario were involuntarily produced. Rather than discarding E2, we chose to proceed with it to test the robustness of our approach against such deviations. Comparing E1 and E2 is difficult because of different time ranges: [305min, 600min] in case of E1 and [158.5min, 220min] for E2. For E2 Tab.2 shows for Δt = 20min, t_1_ = 180min, t_2_ = 200min relative errors of ρt_0_ = 2%, ρT_A0_ = −2% and for Δt = 30min, t_1_ = 200min, t_2_ = 230min relative errors of ρt_0_ = 1%, ρT_A0_ = 12%. In Tab.4 we read for Δt = 10min, t_1,1_ = 180min, t_1,N_ = 199min (N = 20) relative errors ρt_0_ = −4%, ρT_A0_ = 3% and for Δt = 10min, t_1,1_ = 200min, t_1,N_ = 209 (N = 10) relative errors of ρt_0_ = 8%, ρT_A0_ = 14%. Given the deviations of the E2 scenario from the ideal ‘constant-fast-drop-constant’ scenario the overall accurateness of the results in E2 seems not significantly worse than in E1.

Another more practical challenge involves positioning the temperature probes for T_X_ and T_Y_ inside the boxes without opening them, which would destroy the temperature information about T_A0_ in the closed box almost immediately. Here merely experimental methods may lead to results, such as drilling fine holes in cupboard or cardboard walls to insert the probes for T_Z_ measurements. It might even be necessary to introduce a thin optical fiber to explore the arrangement of the boxes’ content before implementing a temperature probe.

Further experiments and reconstructions are necessary to determine the limitations of our approach and to develop methods to address its problems e.g. by using more complex cooling models for boxes in changing T_A_(t). In extreme cases, a finite element simulation of the cooling box could be used to handle strongly varying ambient temperatures. Considering the Newtonian cooling assumption as a first step which could be improved by more complex model assumptions, we believe that deviations of t_0_ - and T_A0_ estimations and of the reconstructed cooling curves (see Fig.2) especially in the phase of short time distances t_1_ - t_0_ could be diminished. Moreover the quadruple measurement distance Δt needs to be chosen carefully. While a too small Δt increases susceptibility to measurement errors, a too large value moves the second measurement into the asymptotic regime of T_Z_(t) converging to T_A1._ The corresponding loss of information would again lead to strongly increasing estimation errors in t_0_^ and T_A0_^. Consequently it would make sense to bound the time interval, where quadruples can be advantageously used for t_0_* and T_A0_* which may be called localization.

The impact of variations in T_A0_ - and t_0_ estimation on TTDE results can be quantified, in principle, (see e.g. [Weiser 2018] for FEM-TTDE) if the TTDE method allows for changing T_A_. Here the t_0_-error *δ*t_0_ plays a special role since it simply shifts the correct TDE-result t_D_^+^ to t_D_^+^+ *δ*t_0_. Even errors of high relative values of 10% or 20% seem to be tolerable if the value *δ*t_0_ stays smaller than the standard deviation of the TDE result. However a valid estimation of the resulting error in TDE from t_0_*- and T_A0_* error is a desideratum. Resulting errors for TDE can be estimated via Gaussian error propagation or Monte Carlo methods as in [Hubig 2010], [Weiser 2018].

In cases of large temperature differences T_A0_ – T_A1_ the usual tolerance interval of MHH is insufficient to handle the resulting TTDE error (see e.g. [Hubig 2010]), which prevents approximate solutions with estimated constant mean ambient temperatures. To address this problem Althaus and Henßge [Althaus 1999] attempted to adapt the MHH model with instantaneous ambient temperature changes but assumed known values for t_0_ and T_A0_. Their approach used a sequential twofold MHH application with an altered MHH model for the first step. The authors fitted the parameter A’s value for their model using experimental dummy cooling data in a scenario with an instantaneous temperature drop from T_A0_ down to T_A1_. Said fitting result for A therefore depends on the temperatures T_A0_ and T_A1_. An error estimation for the resulting TDE could be performed thus via Monte Carlo Simulation in a laborious process.

While our results demonstrate the potential of the method for t_0_ - and T_A0_ reconstruction in principle and suggest its applicability in case work, particularly when the temperature drop occurred recently and moderate variations in TDE are acceptable, further research is needed to explore more elaborate models, estimation techniques or localization algorithms to enhance the methods accuracy. Moreover an estimation of the (t_0_, T_A0_)-reconstruction errors impact on the final TDE error is essential.

## Acknowledgements

The present study is part of a project funded by the Deutsche Forschungsgemeinschaft DFG with the reference numbers MA 2501/4-2 (Prof. Dr. Gita Mall) and WE 2937/10-2 (Dr. Martin Weiser).

## Statements and Declarations

### Compliance with Ethical Standards

No living or deceased humans nor living or dead animals were directly involved in the study.

### Author contributions

Study design and milestone discussions: All authors. Experiments: J. Shanmugam, S. Schenkl, S. Springer. Programming: J. Shanmugam, M. Hubig, M. Weiser. Debugging: J. Sudau, F. Shah. Output interpretation/presentation: J. Shanmugam, M. Hubig, H. Muggenthaler. Paper written/corrected: J. Shanmugam, M. Hubig, M. Weiser, H. Muggenthaler. Paper reading / corrections: All authors.

### Material and/or code availability

The program code is available from the corresponding author on reasonable request.

